# Mythical and Observable Trends in Human Sex Ratio at Birth

**DOI:** 10.1101/2020.04.21.054445

**Authors:** Yanan Long, Qi Chen, Henrik Larsson, Andrey Rzhetsky

## Abstract

The human sex ratio at birth (SRB) is defined as the ratio between the number of newborn boys to the total number of newborns per time unit. It is, typically, slightly greater than 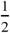 (more boys than girls) and fluctuates over time. In this study, we sought to “myth-check” previously reported associations (and test new hypotheses) using variants of mixed-effect regression analyses and time-series models on two very large electronic health record datasets, representing the populations in the United States and Sweden, respectively. Our results revealed that neither dataset supported models in which the SRB changed seasonally or in response to variations in ambient temperature, and that an increased level of a diverse array of pollutants were associated with lower SRBs. Moreover, we found that increased levels of industrial and agricultural activity, which served as proxies for water pollution, were also associated with lower SRBs.

## Introduction

Because human male gametes bearing X or Y chromosomes are equally frequent (being produced by meiosis symmetrically partitioning two sex chromosomes), and because female gametes bear only X chromosomes, one would expect that the SRB value at conception is exactly 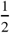. A recent study (***Orzack et al., 2015***) using the fluorescent *in situ* hybridization technique to determine the sex of embryos conceived through *in vitro* fertilization estimated that sex ratio at conception is indeed statistically indistinguishable from one half, 0.502 (***Orzack et al., 2015***). Nonetheless, historically observed SRB values have been, typically, significantly different from 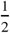 (***Bruckner and Catalano, 2018***), which means that SRBs must fluctuate due to the differential survival of each sex’s embryos and fetuses (***Bruckner and Catalano, 2018***). One possible mechanism for higher SRBs is that more female pregnancies terminate before they are clinically detectable. These excess female losses occur primarily during the first and early-second trimesters, whereas excess male losses occur during the late-second and third trimesters (***Bruckner and Catalano, 2018***). In the late-second and third trimesters, SRB again reduces due to that male-biased mortality.

Previous literature suggests that the SRB can fluctuate with time (***Bruckner and Catalano, 2018***) and is putatively associated with a number of environmental factors, such as chemical pollution, catastrophic events that exert stress on pregnant women (*e.g.*, terrorist attacks and earthquakes), radiation, weather parameters, and even seasons of conception (Table 1). While there are multiple studies in which the positive associations between air pollution and spontaneous abortion have been observed (***Leiser et al., 2019***; ***Zhang et al., 2019***), most of those conclusions were drawn based on analyses of relatively small samples (Table 1). In this study, we harnessed the power of two very large datasets: MarketScan’s insurance claim data (***Hansen, 2017***) in the United States (more than 100 million unique patients and more than three million unique newborns recorded between 2003 to 2011), and Sweden’s birth registry data (∼ 3.2 million newborns from 1983 to 2013) (***Emilsson et al., 2015***). This is the first systematic investigation of numerous chemical pollutants and other environmental factors using large datasets from two continents.

**Table 1.** Exogenous factors reported in the literature to have an impact on the SRB (***Terrell et al., 2011***; ***James and Grech, 2017***). A “-” indicates that sample sizes were not mentioned in the articles reporting or reviewing the corresponding results.

| Exogenous Factor | Number of Studies | Sample Size |
| --- | --- | --- |
| Dioxins (*Terrell et al., 2011*) | 13 | 291 |
| Polychlorinated biphenyls (PCBs) (*Terrell et al., 2011*) | 9 | 98 |
| 1,2-Dibromo-3-chloropropane (DBCP) (*Terrell et al., 2011*) | 2 | 29 |
| Dichlorodiphenyltrichloroethane (DDT) (*Terrell et al., 2011*) | 4 | 1623 |
| Hexachlorobenzene (HCB) (*Terrell et al., 2011*) | 2 | 262 |
| Vinclozolin (*Terrell et al., 2011*) | 1 | 95 |
| Multiple pesticides (*Terrell et al., 2011*) | 5 | 382 |
| Lead (*Terrell et al., 2011*) | 5 | 6566 |
| Methylmercury (*Terrell et al., 2011*) | 1 | 4808 |
| Multiple metals (*Terrell et al., 2011*) | 10 | 1015 |
| Non-ionizing radiation (*Terrell et al., 2011*) | 12 | 2926 |
| Ionizing radiation (*Terrell et al., 2011*) | 15 | 4959 |
| Seasonality (*Lerchl, 1998; Melnikov and Grech, 2003*) | 2 | - |
| Ambient temperature (*Catalano et al., 2008; Helle et al., 2009; Fukuda et al., 2014; Dixon et al., 2011*) | 4 | - |
| Economic stress (*Catalano, 2003*) | 1 | - |
| Terrorist attacks (*James and Grech, 2017*) | 2 | - |

## Methods

### Data

The IBM Health MarketScan data (***Hansen, 2017***) represents 104, 565, 671 unique individuals and 3, 134, 062 unique live births. The Swedish Health Registries (***Emilsson et al., 2015***) record health statistics for over ten million individuals, and 3, 260, 304 unique live births. We juxtaposed time-stamped birth events in the two countries with exogenous factor measurements retrieved from the US National Oceanic and Atmospheric Administration, the US Environmental Protection Agency, the Swedish Meteorological and Hydrological Institute, and Statistics Sweden. We used a subset of the MarketScan data (***Hansen, 2017***) which contained information about live births during the period of 2003 to 2011 with county information encoded in FIPS (Federal Information Processing Standards) codes and a family link profile indicating the composition of the families contained in the dataset. The date, geographic distribution, and the mothers of the newborns can be directly extracted from these datasets. For environmental factors, we used the Environmental Quality Index (EQI) data compiled by the United States Environmental Protection Agency (EPA) (***Lobdell et al., 2011***; ***Messer et al., 2014***).

### Cluster Analysis

In order to simplify subsequent analyses, we first reduced the dimensionality of the EQI dataset by performing hierarchical clustering analysis on the matrix of Spearman’s rank correlation coefficients (*ρ*) with the Ward’s method. We then used the R-language (***R Core Team, 2017***) package `pvclust` (***Suzuki and Shimodaira, 2006***) to minimize the total within-cluster variance (***Legendre and Legendre, 2012***). The resulting dendrogram and list of factors can be found in the SI. Each cluster contains at least two factors and is represented by the mean of all the elements in the cluster.

### Regression Analysis

We used Bayesian logistic regression with random effects, implemented in the R-language package `brms` (***Bürkner, 2017***) and the No-U-Turn sampler (NUTS) (***Hoffman and Gelman, 2014***). The model for the *j*^th^ factor (predictor) is given as follows:

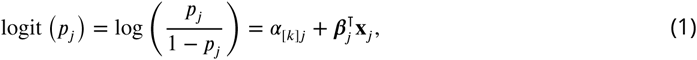

where *p*_*j*_ is the probability that a newborn is male, **x**_*j*_ is the vector representing the *j*^th^ factor, ***β***_*j*_ their coefficients, and *α*_[*k*]*j*_ the intercept for the *k*^th^ group-level, representing states or counties in the US, and *kommuner* (municipalities) or *län* (counties) in Sweden, if applicable. The group-level effect is modeled for a single random effect by (***Gelman and Hill, 2006***)

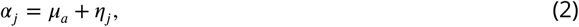

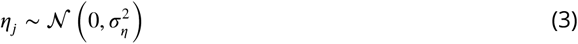

and for two random effects, representing e.g. state- and county-specific effects, by

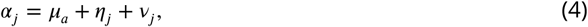

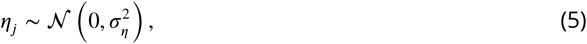

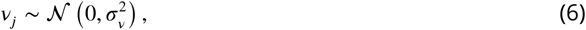

where *η*_*j*_ and *v*_*j*_ are independent of each other and for all *j*. Moreover, we partitioned the independent variables into septiles, so that ***β***_*j*_ ∈ ℝ^6^, with one regression coefficient for each of the six septiles other than the first, which was chosen as the reference level (***Khan et al., 2019***).

We applied logistic regression in two ways. First, to assess the effect of environmental factors, we regressed the septiles of each of the individual factors against the SRB, with each sample point representing a county. Therefore, each septile, aside from the baseline, has a coefficient. Second, to test whether a mother’s diagnostic history (DX) affected the SRB, we regressed a DX’s indicator variables against the SRB, with each sample point representing a newborn/mother pair. For model selection in both cases, we performed repeated (10 times) 10-fold cross-validation and calculated the information criterion relative to the null model (where **x**_*j*_ = **0**, i.e., the model comprised solely of the intercept). We computed the average difference in information criterion (ΔIC) and standard error (SE) for each factor, and used the Benjamini–Yekutieli method to account for multiple comparisons (***Benjamini et al., 2001***).

### Univariate Time-series Analysis

To assess the effect of one-off, stressful events on the SRB, we used two different time series techniques. We first fitted seasonal univariate autoregressive, integrated, moving average (sARIMA) models using the Box-Jenkins method (***Box et al., 2015***) in conjunction with monthly (28-day periods) and weekly live birth data up to the event and then performed an out-of-sample prediction. A sARIMA model is given as follows:

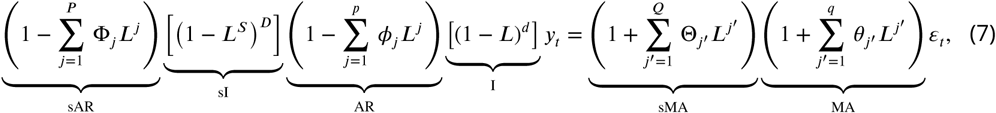

where AR indicates the auto-regression term, I the integration term, and MA the moving average term (an s before any of the above stands for seasonal), *y*_*t*_ the observed univariate time series of interest, ***L*** the lag operator such that ***L***(*y*_*t*_) = *y*_*t*−1_, *ε*’s white noises, *s* ⩾ 2 the degree of seasonality (i.e., the number of seasonal terms per year), and *ϕ*’s, *θ*’s, Φ’s and Θ’s are model parameters to be estimated. We used the R package’s `forecast` (***Hyndman and Khandakar, 2008***; ***Hyndman et al., 2019***) `auto.arima` function to fit the data, which performs a step-wise search on the (*p, d, q*, ***P***, ***D, Q***) hyperparameter space and compares different models by using the Bayesian Information Criterion (BIC) (***Schwarz, 1978***). We confirmed the optimal models’ goodness-of-fit using the Breusch-Godfrey test on the residuals, which tests for the presence of autocorrelation up to degree *s* (***Breusch, 1978***; ***Godfrey, 1978***; ***Hayashi, 2000***). On the other hand, we fitted the same data as above to Bayesian structural time series (BSTS) models, which are state space models given in the general form by (***Scott and Varian, 2013***):

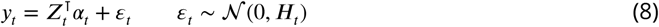

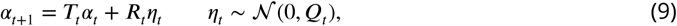

where *y*_*t*_ is the observed time series and *α*_*t*_ the *unobserved* latent state. In particular, we used the local linear trend model with additional seasonal terms (***Murphy, 2012***; ***Scott and Varian, 2013***):

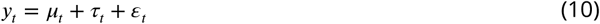

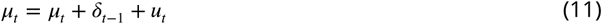

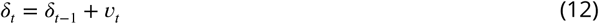

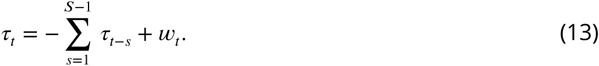

Under this formulation, we have *η*_*t*_ = [*u*_*t*_ *v*_*t*_ *w*_*t*_] and ***Q***_*t*_ is a *t*-invariant block diagonal matrix with diagonal elements 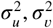 and 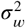. Furthermore, we write *α*_*t*_ = [*µ*_*t*_ *δ*_*t*_ *τ*_*t*−*S*+2_ … *τ*_*t*_], which implies that both ***Z***_*t*_ and ***T***_*t*_ are *t*-invariant matrices of 0’s and 1’s such that Equations 10–13 hold. We used the R package `CausalImpact` (***Brodersen et al., 2015***), which uses the R package `bsts` (***Scott, 2019***) as backend, to fit the data.

### Correlation and Causality

To test whether the SRB was effected by ambient temperature, we grouped daily SRB data and temperatures into 91-day (13-week) periods and calculated the Pearson correlation coefficient (*r*) between each SRB and ambient temperature. We then performed the Student’s *t*-test for the null hypothesis that the true correlation is 0. Furthermore, we fitted the SRB/temperature pair to a vector autoregression (VAR) model for a maximum lag order of 4 (52 weeks), using the BIC as the metric for model selection, and then tested for the null hypothesis of the non-existence of Granger causality using the ***F*** -test (***Granger, 1969***).

## Results

We start by describing the negative results concordant across the two datasets. Our model selection results rejected the whole spectrum of models that postulate periodic, annual SRB changes (***Lerchl, 1998***; ***Melnikov and Grech, 2003***). For both US and Swedish datasets, the best-fitting sARIMA model described the SRB as lacking seasonality throughout the year. Similarly, when we tested the claim that ambient temperatures during conception affect the SRB (***Catalano et al., 2008***; ***Helle et al., 2009***; ***Fukuda et al., 2014***; ***Dixson et al., 2011***), we found that neither dataset supported this association, as both the Student’s *t*-test and the ***F*** -test concluded that the SRB was independent of ambient temperature measurements (Table S7).

A comparison of each dataset’s environmental measurements revealed that Sweden enjoyed both lower variations and lower mean values of measured concentrations of substances in the air. Unfortunately, the Swedish dataset also provided fewer measured pollutants, which made our cross-country analysis more difficult. Figure 1**A** shows a comparison of pollutant concentration distributions in both countries. The US environmental measurements dataset presented its own difficulty: Many pollutants appeared highly collinear in their spatial variation. To circumvent this, we performed a cluster analysis on the environmental factors, subdividing them into 26 clusters (see the *Cluster Analysis* section in *Methods* and Table 4). All pollutants within the same cluster were highly correlated, while the correlation between distinct clusters was smaller, allowing for useful inferences of associations between SRB changes and environmental states.

**Figure 1.**
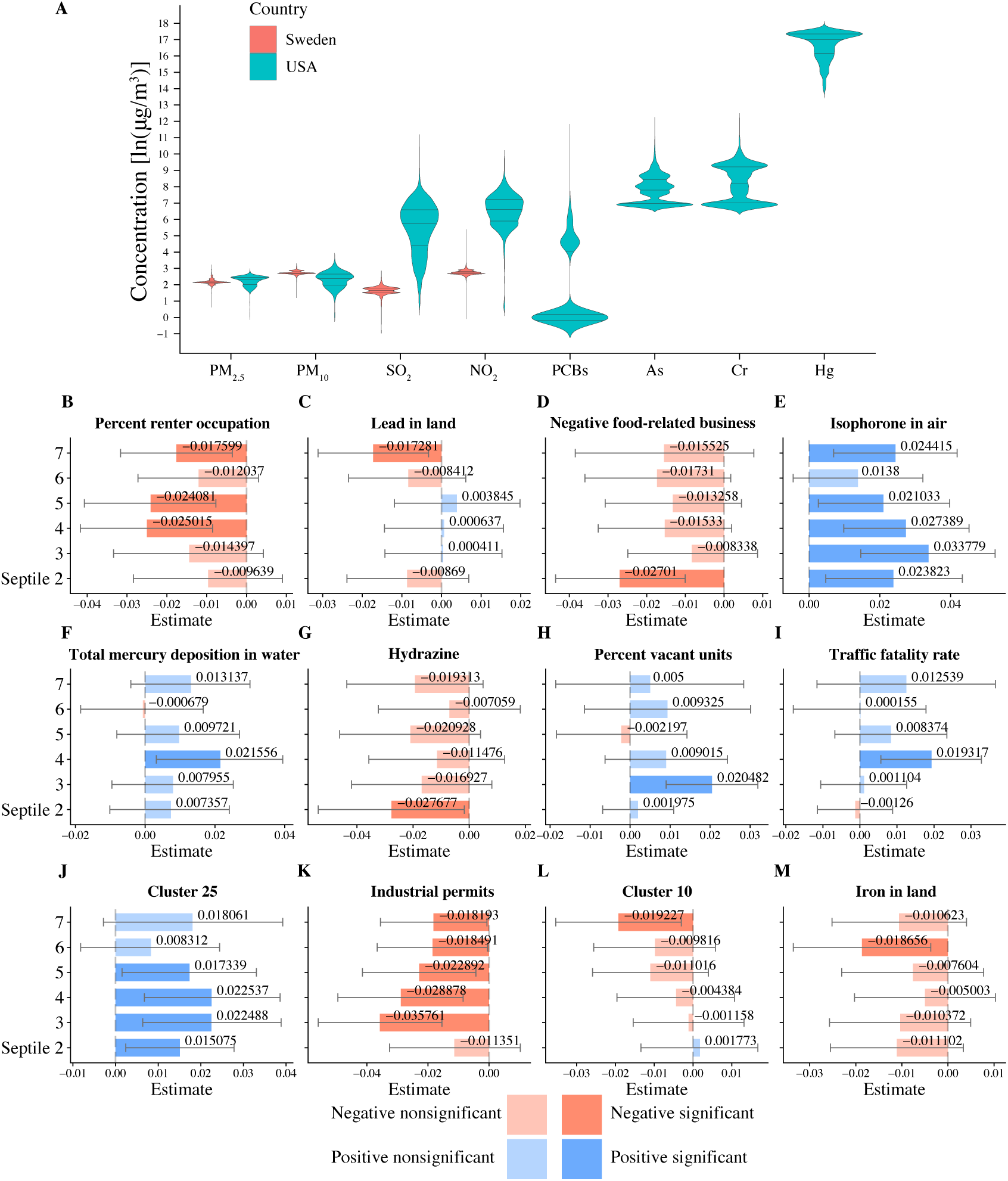
Airborne health-related substances and their association with the SRB. **A** Comparison of airborne pollutant concentrations across the US (cyan violin plots) and Sweden (pink violin plots). Only four air components, fine particulate matter (PM_2.5_), coarse particulate matter (PM_10_), sulfur dioxide (SO_2_), and nitrogen dioxide (NO_2_) are measured in both countries. US counties appear to have higher mean pollution levels and are more variable in terms of pollution. **B-M**. A sample of 12 one-environmental factor logistic regression models that are most explanatory with respect to SRB. For each environmental factor, we partitioned counties into seven, equal-sized groups (septiles), ordered by levels of measurements, so that septile one corresponds to the lowest and septile seven to the highest concentration. Each plot shows bar plots of regression coefficients and 95%confidence intervals (error bar) of the second to the seventh septiles, with the first septile chosen as the reference level. The 12 models were ranked by the statistically significant factor’s strength of association with at least one statistically significant coefficient by decreasing ΔIC; septiles whose coefficients are not significantly different from 0 at the 95% confidence level were plotted with a reduced alpha level. Blue bars represent positive coefficients, whereas red bars represent negative coefficients. “Negative food-related businesses” is a term used by the Environmental Protection Agency’s Environmental Quality Index team and is explained as “businesses like fast-food restaurants, convenience stores, and pretzel trucks.” “Percent vacant units” stands for “percent of vacant housing units.” Substances contributing to cluster ten and cluster 25 are listed in Table 4. See Table S11 for more details regarding the factors’ and clusters’ identities.

Using the US dataset, we were able to validate a number of previous studies’ findings regarding the association between the SRB and exogenous factors (Table 2). Specifically, our data suggests that PCBs (polychlorinated biphenyls), aluminium in air, chromium in water, and total mercury appear to drive the SRB up, while lead in soil appears to be associated with a decreased SRB. Meanwhile, we refute a number of previous reports (these are indicated with a dash in the second column in Table 2). Finally, we also found a number of new environmental associations that have not been reported before (see Table 3, Figure 1 **B–M**, and Figure 1–figure supplement 1). Figure 1 **B–M** shows that increased pollutant levels appear to be associated with both increased and decreased SRB values (Plates **E**,**F**,**H**,**I**, and **J**, and the remaining Plates, respectively).

**Table 2.** Test results for factors selected from the literature reports (Table 1). A factor was included only if both its ΔIC and the coefficient of at least one of its septiles were statistically significant.

| Factor name | effect |
| --- | --- |
| PCBs (air and water) | ↑ |
| DBCP (water) | — |
| Lead (land) | ↓ |
| Lead (air) | — |
| Aluminium (air) | ↑ |
| Chromium (air) | — |
| Chromium (water) | ↑ |
| Arsenic (land) | — |
| Arsenic (water) | ↑ |
| Cadmium (air and water) | — |
| Total mercury deposition | ↑ |
| Violent crime rate | — |
| Unemployed rate | — |
| Working out of county (long commute) | — |

**Table 3.** Test results for additional factors with statistically significant effects. A factor was included only if both its ΔIC and the coefficient of at least one of its septiles were statistically significant.

| Factor name | effect |
| --- | --- |
| Iron | ↓ |
| Nitrate | ↑ |
| 2-Nitropropane | ↑ |
| Carbon monoxide | ↑ |
| Bis-2-ethylhexyl phthalate | ↓ |
| Ethyl chloride | ↑ |
| Isophorone | ↑ |
| Hydrazine | ↓ |
| Phosphorus | ↑ |
| Quinonline | ↓ |
| Extreme drought | ↑ |
| Traffic fatality rate | ↑ |
| Industrial permits per 1000 km of stream | ↓ |
| Animal units | ↓ |
| Irrigation | ↓ |
| Negative food related businesses | ↓ |
| Renter occupation | ↓ |
| Vacant units | ↑ |

**Table 4.**
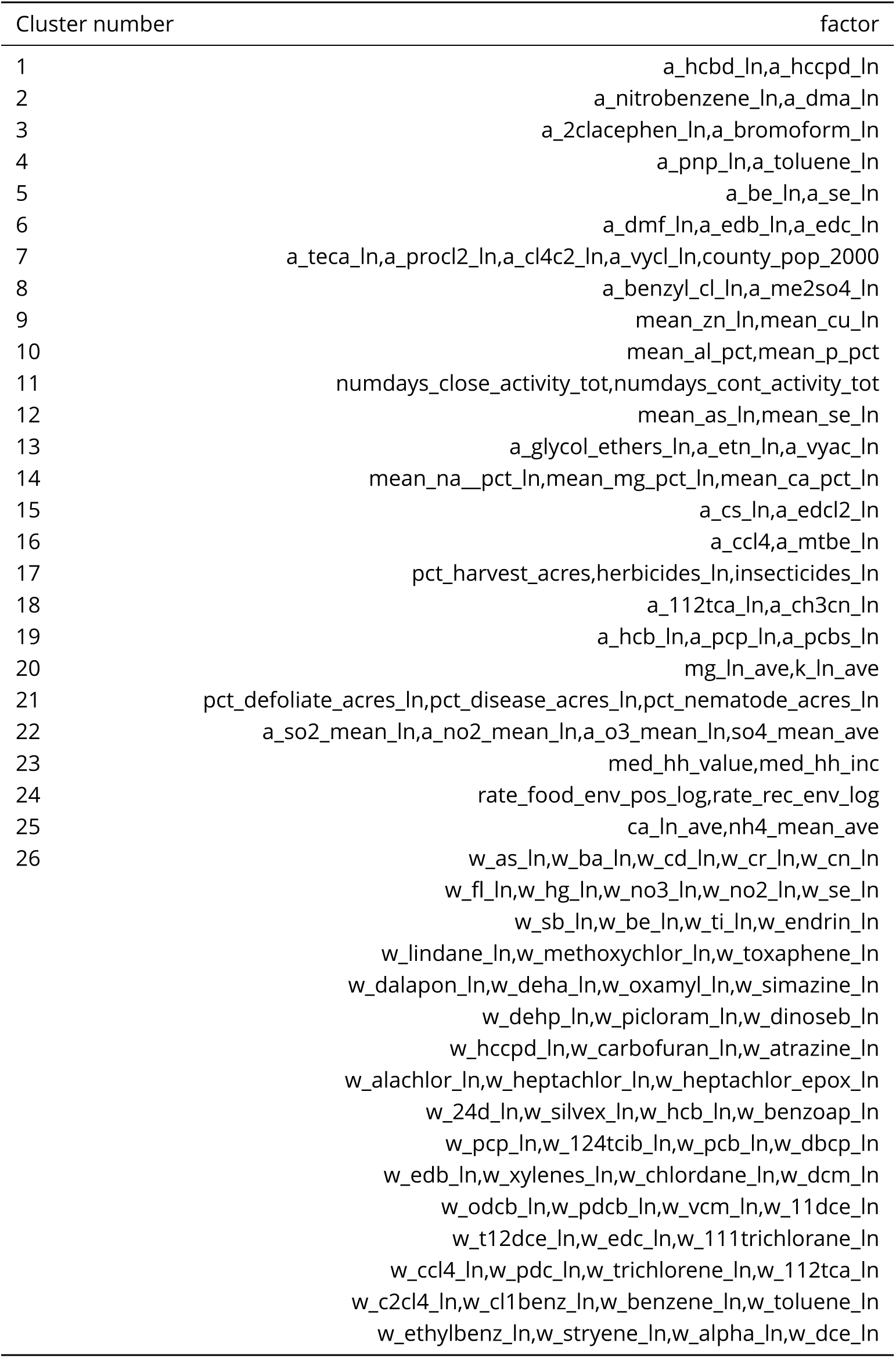
Clusters obtained by applying the Ward’s method to the EQI dataset

**Table 5.** List of the top 8 statistically significant environmental factors or clusters of factors with random effects at the county level from EQI by ΔIC. For the full table, see Table S3

| Factor | explanation | ΔIC | SE |
| --- | --- | --- | --- |
| pct_rent_occ | percent renter occupation | 54.2185 | 8.7760 |
| cluster_ward_9 | zinc (Zn) and copper (Cu) in air | 52.3907 | 8.8256 |
| mean_pb_ln | lead (Pb) in land | 51.7339 | 9.6749 |
| farms_per_acre_ln | farms per acre | 51.6266 | 9.0878 |
| cluster_ward_15 | caesium (Cs) and ethylidene dichloride ($\text{CHCl}_2\text{CHCl}_2$) in air | 51.5511 | 8.5623 |
| pct_mt_10units_log | percent housing with $\geq 10$ units | 51.3259 | 8.5483 |
| rate_food_env_neg | negative food-related businesses (e.g. junk food) | 51.3230 | 9.2907 |
| a_isophorone_ln | isophorone in air | 50.9430 | 10.3995 |

The geographic distribution of these pollutants varies remarkably, as seen in Figure 1–figure supplement 1. For example, lead in land (Figure 1–figure supplement 1**B**) appears to be enriched in the northeast, southwest, and mid-east US, but not in the south. Hydrazine (Figure 1–figure supplement 1**F**) appears to follow capricious, blotch-like shapes in the eastern US, each blotch likely centered at a factory emitting this pollutant. Total mercury deposition in water (Figure 1–figure supplement 1**E**) mostly affects eastern US states with the heaviest load in the northeastern states. It is this variability in the distribution of various substances in the environment that makes teasing out individual associations possible.

Finally, when we tested links between two stressful events in the US (Hurricane Katrina and the Virginia Tech shooting) and the SRB using seasonal autoregressive integrated moving-average (sARIMA) models and state-space models (SSMs) (see the *Univariate time-series analysis* section in *Methods*), we were able to identify significant associations only in the case of the Virginia Tech shooting – the SRB was lower than expected 34 weeks after the event (see Figures 2c and 3c as well as Tables S5(c) and S6(c)).

**Figure 2.**
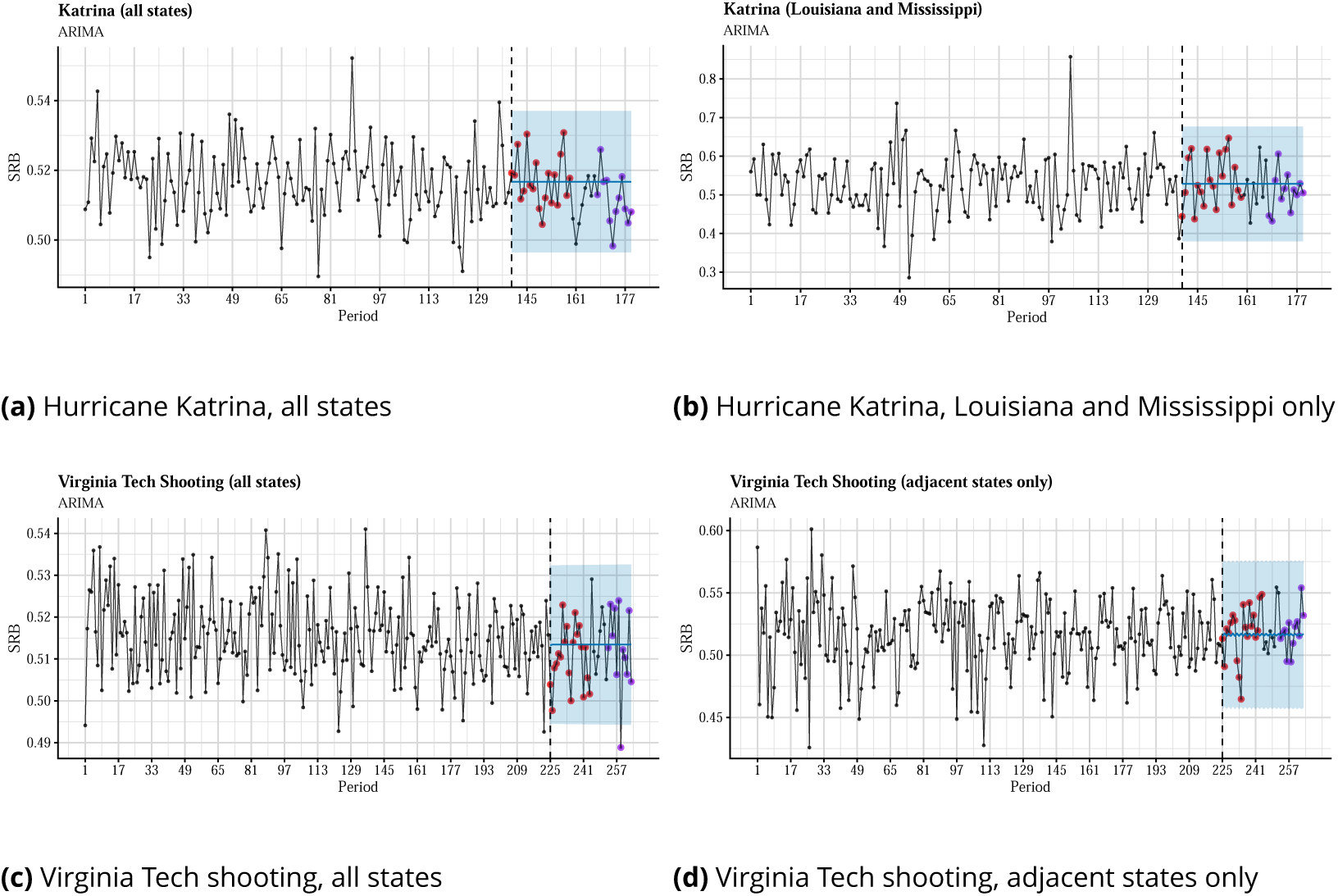
Time series plots and out-of-sample forecasts for SRB data grouped into 7-day periods and fitted with seasonal ARIMA models. The blue shade is the 95% confidence level. The observed SRBs for the first 5 months after the intervention are presented by red dots, whereas the observed SRBs for 7–9 months after the intervention are presented by purple dots.

**Figure 3.**
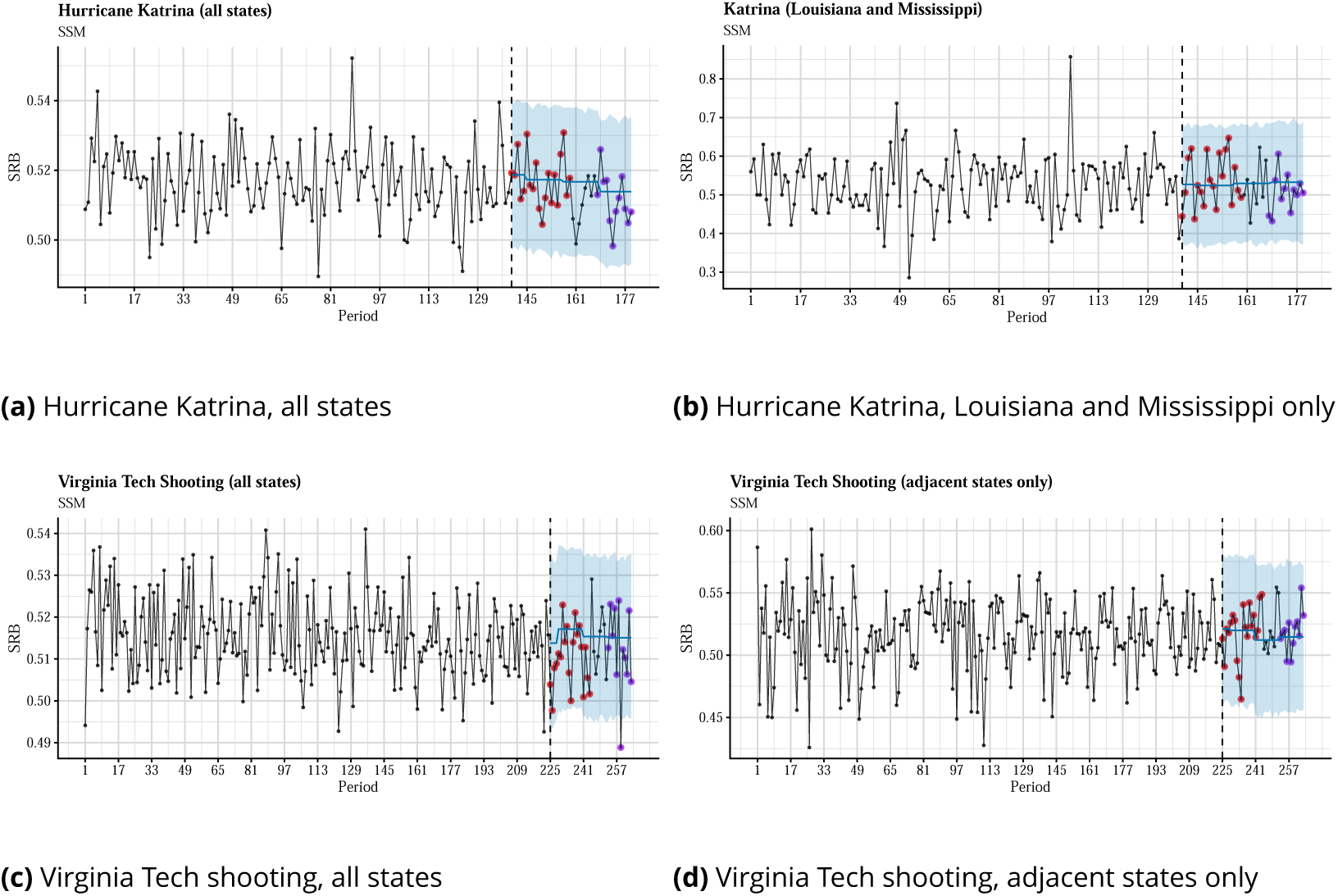
Time series plots and out-of-sample forecasts for SRB data grouped into 7-day periods and fitted with state-space models. The blue shade is the 95% confidence level. The observed SRBs for the first 5 months after the intervention are presented by red dots, whereas the observed SRBs for 7–9 months after the intervention are presented by purple dots. **(c)** Virginia Tech shooting, all states **(d)** Virginia Tech shooting, adjacent states only

## Discussion

The literature is replete with theories striving to explain SRB changes in terms of evolutionary stresses and benefits. For example, one such theory, named the Trivers-Willard hypothesis after the researchers who proposed it (***Trivers and Willard, 1973***), states that the SRB fluctuates due to environmental conditions changing from favourable to unfavourable and back. Under favourable conditions, a male descendent is postulated to generate many more offspring than a female, and vice versa under unfavourable circumstances. Under this theory, parents maximize the number of grandchildren, thereby regulating the SRB to this end. The empirical support for this hypothesis is controversial (***Keller et al., 2001***; ***Lynch et al., 2018***; ***Song, 2018***), and the theory is vague when it comes to elucidating the underlying biological mechanisms. Taken as whole, our results appear to cast serious doubt on the validity of the Trivers-Willard hypothesis.

Because the average observed SRB is higher than 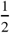, we propose to explain these observations through the differential survival of either gametes or fetuses. Specifically, we hypothesize that female fetuses are subject to a higher risk of termination than male ones, which seems contrary to the theory that the SRB skews towards males (***Orzack et al., 2015***). To explain this inconsistency, some researchers have suggested that more female than male pregnancies terminate before they are clinically detectable, that is, excess female losses occur primarily during the first and early-second trimesters, whereas excess male losses occur during the late-second and third trimesters (***DiPietro and Voegtline, 2017***; ***Bruckner and Catalano, 2018***). In other words, the SRB starts at very close to ^1^, then increases due to female-biased mortality, and finally reduces due to male-biased mortality. Because the physiology of male and female fetuses differs (***DiPietro and Voegtline, 2017***), it is conceivable that specific environmental stresses affect fetuses of different sexes in a sex-specific way.

Recent studies have uncovered a positive association between pollution and spontaneous abortion (***Leiser et al., 2019***; ***Zhang et al., 2019***), so our analyses presented here allow for more specific hypotheses regarding the potential chemicals that might cause preferential abortions of male or female fetuses. Fortunately, most of the putative positive associations identified in our study are experimentally testable, at least with animal models. However, given the large sample sizes of our analysis, one would expect that it is rather unlikely that the associations rejected in our study, such as those between SRB and ambient temperature and seasonality, as well as some environmental factors (see Tables 2 and 3), will be found as valid in other large datasets or laboratory studies.

Because the observed, statistically-significant SRB fluctuations have to be mediated by undetected spontaneous abortions, it is important to better understand the underlying mechanisms. The current literature is regretfully meagre regarding causal mechanisms behind the temporal fluctuations of the SRB as well as the plausible ramifications of the presence of environmental toxins and their possible remedies.

Finally, our study also highlights the need for more detailed and systematic measurements of environmental quality data – spatially, temporally, and across the chemical spectrum. For the US, the current measurements are available at a very coarse, county-level scale and are not repeated frequently enough. Ideally, the measurements would be taken frequently enough (say, daily or weekly) to enable the application of time-series methods (e.g. Granger causality or cointegration tests), which would help determine the causal structures of the time variation of exogenous factors on the SRB (***Samartsidis et al., 2018***). On the other hand, only a small subset of US environmental quality measurements were available in Sweden. As a result, our Swedish analyses lacks both the statistical power and breadth of measurements required to reproduce the kinds of positive, statistical associations between SRB and individual environmental factors that we discovered in the US data.

## Acknowledgments

We are grateful to E. Gannon and M. Rzhetsky for comments on earlier versions of this manuscript. This work was funded by the DARPA Big Mechanism program under ARO contract W911NF1410333, by National Institutes of Health grants R01HL122712, 1P50MH094267, and U01HL108634-01, and by a gift from Liz and Kent Dauten.

The authors declare that they have no competing financial interests.

---

**Figure 1–figure supplement 1.**
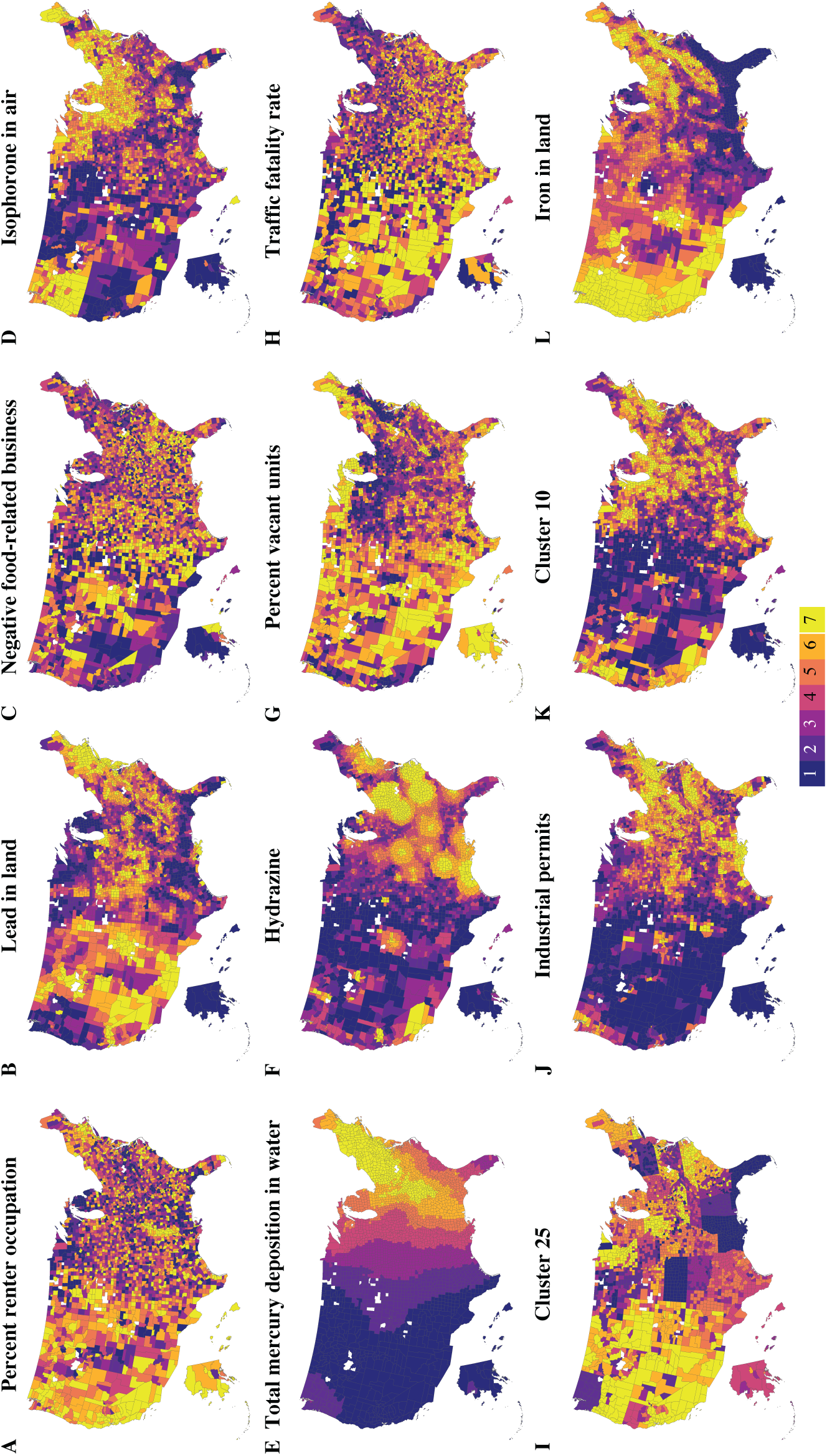
County-level geographical septile distribution for the first 12 statistically significant factors with at least one statistically significant coefficient ranked by decreasing ΔIC. The factors labelled **A–M** are the same as shown in Figure 1, Plates **B–M** and are ordered identically in both figures.

**Figure 1–figure supplement 2.**
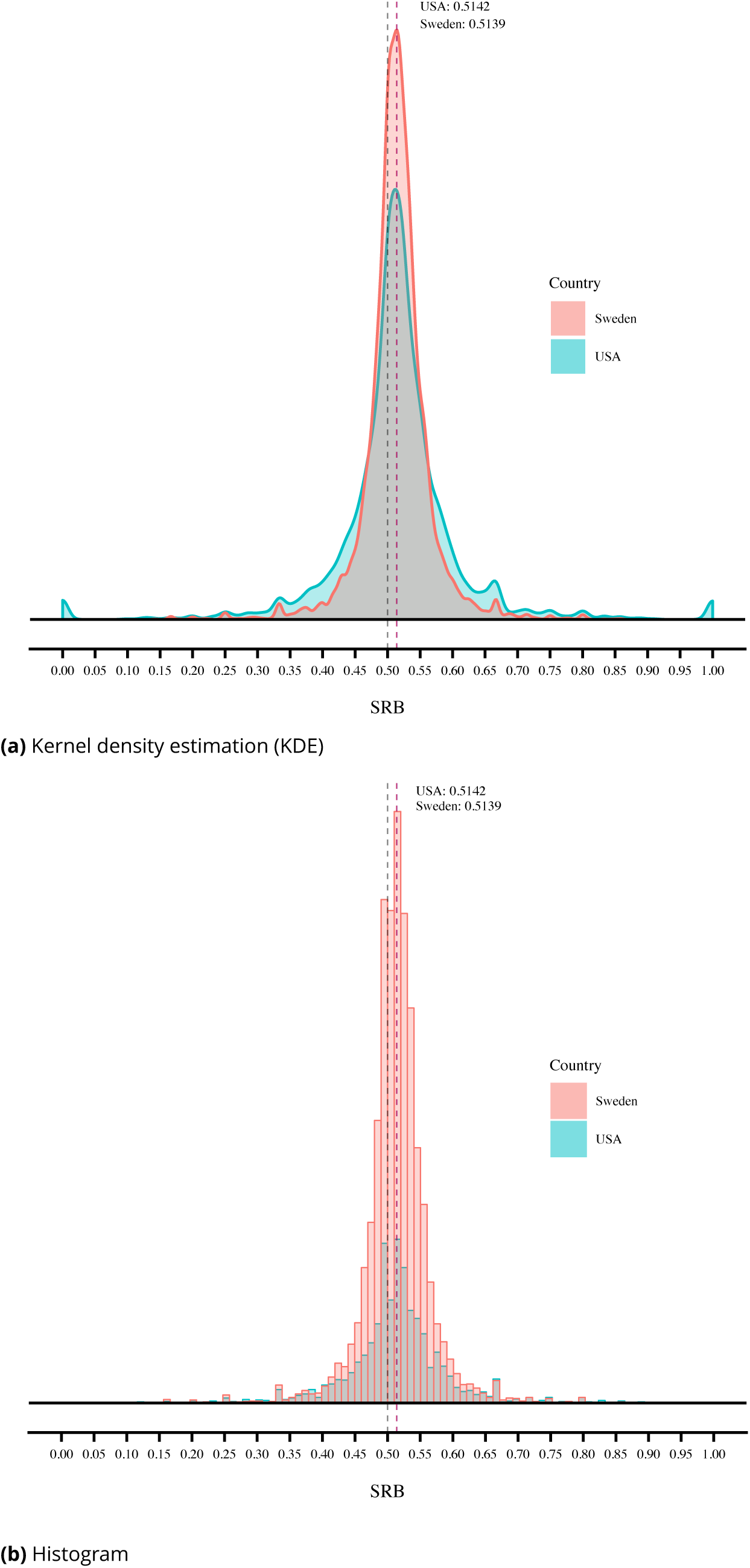
Distribution of the SRB in the US and Sweden at the county level (US) or the kommun level (Sweden) Virginia Tech shooting, all states **(d)** Virginia Tech shooting, adjacent states only

**Figure 1–1gure supplement 3.**
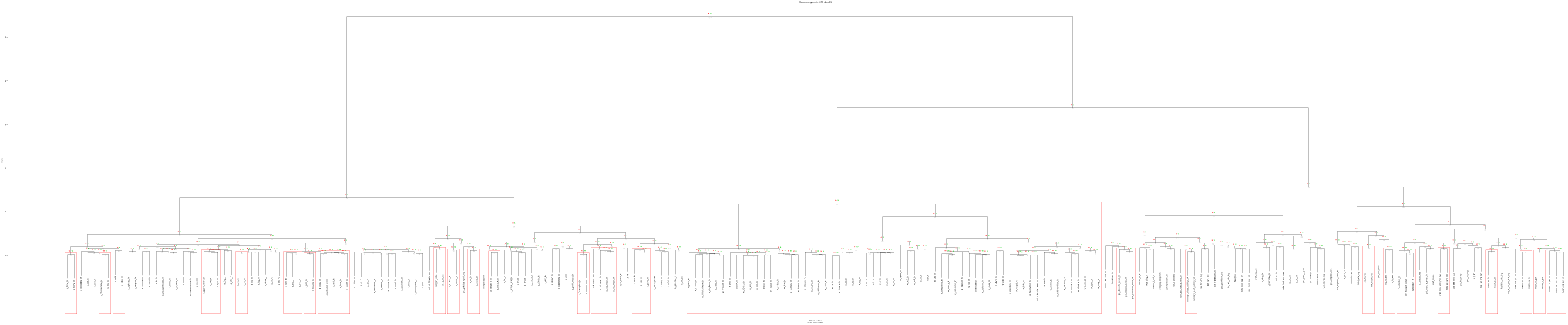
Dendrogram with statistically significant clusters (95% level) in red boxes

**Figure 2–figure supplement 1.**
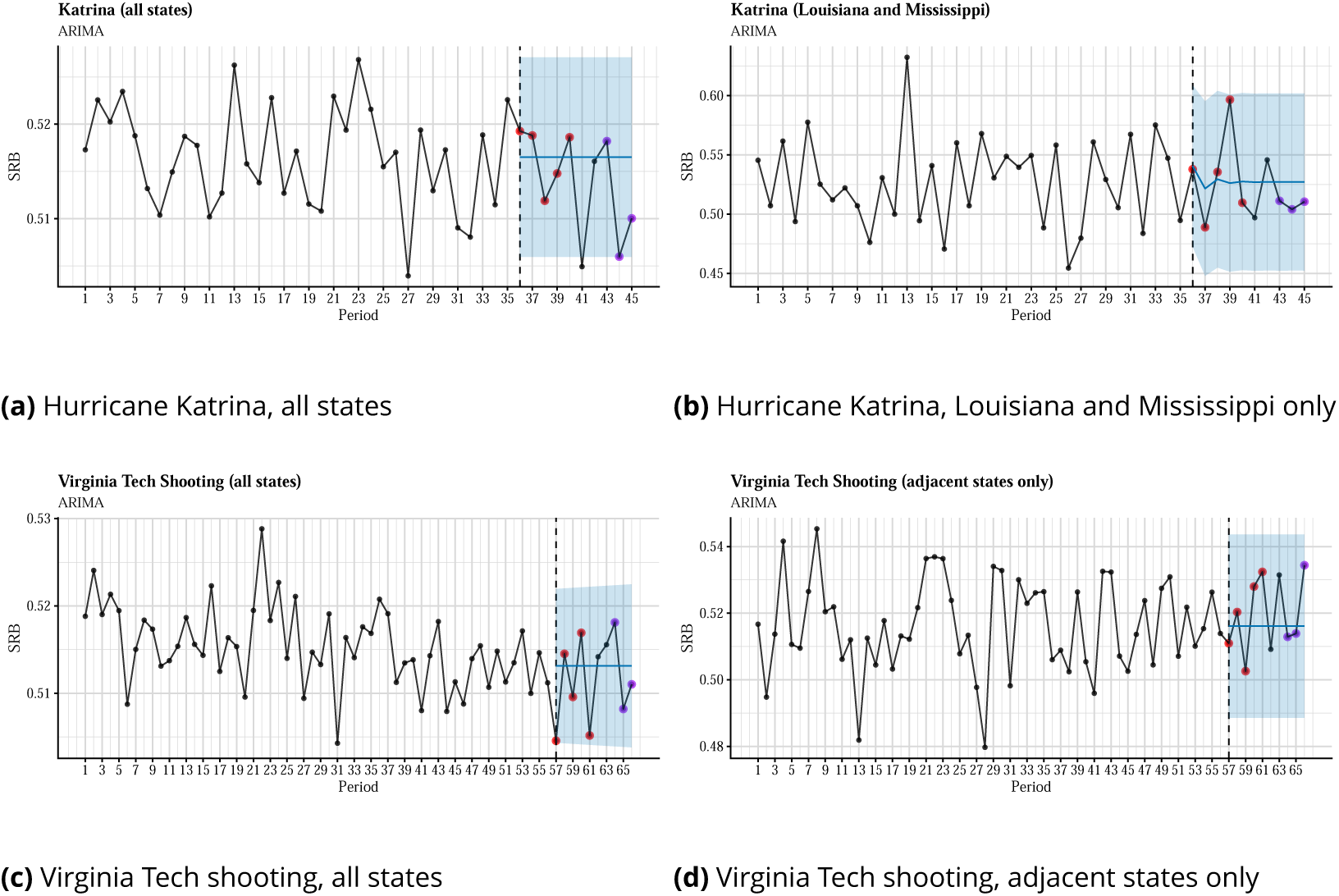
Time series plots and out-of-sample forecasts for SRB data grouped into 28-day periods and fitted with seasonal ARIMA models. The blue shade is the 95% confidence level. The observed SRBs for the first 5 months after the intervention are presented by red dots, whereas the observed SRBs for 7–9 months after the intervention are presented by purple dots.

**Figure 3–figure supplement 1.**
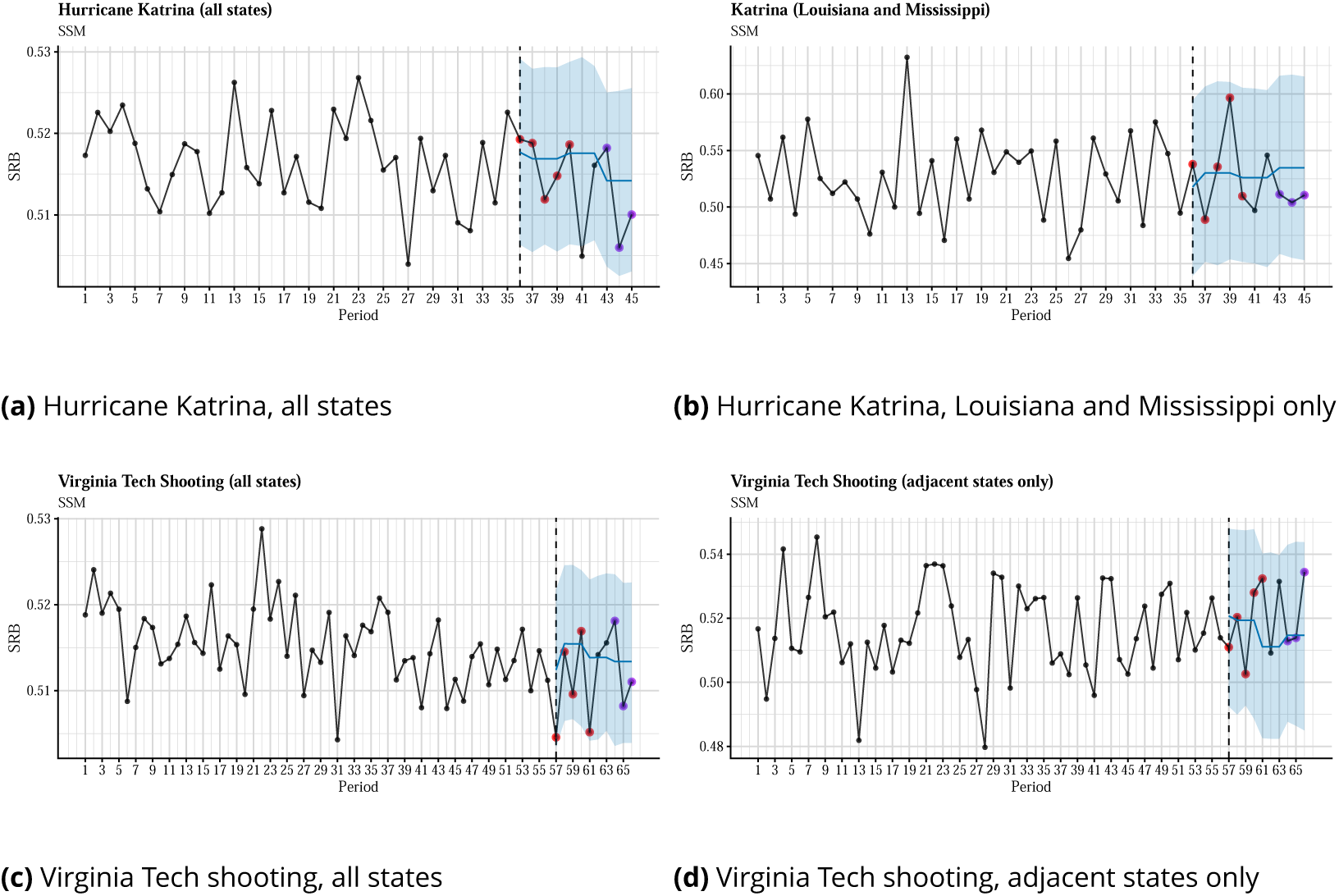
Time series plots and out-of-sample forecasts for SRB data grouped into 28-day periods and fitted with seasonal ARIMA models. The blue shade is the 95% confidence level. The observed SRBs for the first 5 months after the intervention are presented by red dots, whereas the observed SRBs for 7– 9 months after the intervention are presented by purple dots.

